# Fast and interpretable non-negative matrix factorization for atlas-scale single cell data

**DOI:** 10.1101/2021.09.01.458620

**Authors:** Zachary J. DeBruine, J. Andrew Pospisilik, Timothy J. Triche

## Abstract

Non-negative matrix factorization (NMF) is a popular method for analyzing strictly positive data due to its relatively straightforward interpretation. However, NMF has a reputation as a less efficient alternative to the singular value decomposition (SVD), a standard operation that is highly optimized in most linear algebra libraries. Sparse single-cell sequencing assays, now feasible in thousands of subjects and millions of cells, generate data matrices with tens of thousands of strictly non-negative transcript abundance entries. We present an extremely fast NMF implementation made available in the RcppML (Rcpp Machine Learning library) R package that rivals the runtimes of state-of-the-art Singular Value Decomposition (SVD). NMF can now be run quickly on desktop computers to analyze sparse single-cell datasets consisting of hundreds of thousands of cells. Our method improves upon current NMF implementations by introducing a scaling diagonal to increase interpretability, guarantee consistent regularization penalties across different random initializations, and symmetry in symmetric factorizations. We use our method to show how NMF models learned on standard log-normalized count data are interpretable dimensional reductions, describe interpretable patterns of coordinated gene activities, and explain biologically relevant metadata. We believe NMF has the potential to replace PCA in most single-cell analyses, and the presented NMF implementation overcomes previous challenges with long runtime.

## Introduction

In 1999, Lee and Seung proposed an algorithm for Non-Negative Matrix Factorization (NMF) that was able to learn the parts of faces from a set of images (Lee & Seung, 1999). This demonstration that non-negativity constraints enforced an additive decomposition contrasted sharply with other then-popular dimensional reductions such as PCA or SVD which describe axes of variance rather than parts of the whole. Two decades later, a plethora of specialized machine learning algorithms have emerged that incorporate non-negativity constraints, and yet none are more interpretable than a simple non-negative matrix factorization.

Just as a matrix of pixels encoding a face might be decomposed into factors describing the parts of a face, a matrix of the RNA transcript portfolios of single cells may be decomposed into constituent cellular processes (Brunet et al., 2004; Clark et al., 2019). In turn, these processes are orchestrated by coordinated patterns of gene expression. While NMF exposes basic biology and generalizable information amenable to transfer learning (Stein-O’Brien et al., 2019), SVD and PCA capture sequential normalizations or rotations of coordinate axes explaining variation specific to the dataset of interest.

Single-cell count data is strictly non-negative. Non-negative count data is well-suited to non-negative dimension reduction. If the dimensional reduction may contain positive and negative loadings, factors in the learned model could cancel one another out. This happens in SVD and PCA, where each factor gives a linear transformation of the high-dimensional data in low-dimensional space that both over- and under-corrects for all transformations applied by the preceding factors. In contrast, factors in NMF are mutually interdependent and collectively additive, not sequential, and can thus be understood individually. Unfortunately for SVD and PCA, this means it is possible to handle dropout events in both directions – by imputing missing signal (which is good) and by ignoring true signal (which is bad). NMF is not subject to this issue as it cannot contain negative values, thus missing signal is generally imputed up to a point that minimizes mean squared error of reconstruction (Lin & Boutros, 2020).

Many modern approaches to single-cell analysis still rely heavily on variance-based decompositions (SVD or PCA) (Qiu et al., 2017; Stuart et al., 2019). This is because PCA is relatively fast, historically established, well-supported in popular frameworks, and can generate reasonable dimension reductions for visualization and graph-based clustering. Other methods for single-cell analysis use variational autoencoders (Lopez et al., 2018), generative adversarial networks (Marouf et al., 2020), and other new machine learning algorithms, all with some amount of success for their respective objectives. Unfortunately, too many of these algorithms require extensive hyperparameter optimization and thus deep knowledge of the method for proper use, and even if outputs satisfy objectives, most of the learned latent factor models are themselves hardly interpretable.

Given its interpretability, why has NMF not been widely adopted by the single-cell community? In our opinion, the single greatest barrier is computing time. While there are many fast NMF-inspired algorithms for sparse recommender systems, such applications differ significantly from the objectives in single-cell analysis. For example, collaborative filtering and implicit recommenders require different algorithms than simple NMF. To our knowledge, there are no NMF implementations for large sparse matrices that are fast enough to support rapid experimentation, even in high-performance computing environments.

In addition to optimizing runtime, more clear demonstrations of the usefulness of NMF in single-cell analyses are needed. For instance, NMF can serve as both a dimensional reduction in place of PCA (Elyanow et al., 2020) and a resource for learning novel biology (Clark et al., 2019). With sufficient meta-analysis and dataset integration, it is possible that latent factors learned by NMF may replace manually curated toolkits such as GO terms in many gene set enrichment analyses (Stein-O’Brien et al., 2018).

In this manuscript we make the case for using NMF in single-cell analysis rather than PCA. We then present technical aspects of the implementation that have enabled the fast performance of our implementation, which may be of interest to a more interdisciplinary audience. Finally, we direct attention to several perpetual challenges of NMF as a method and propose some avenues for ongoing research.

## Results

### Dimension reduction with NMF is as sensitive as PCA

To make a case for NMF as a sensitive dimensional reduction, we analyzed 103,000 single blood cells from the Human Embryonic Cell Atlas. First, the optimal rank for a factorization was determined by plotting the mean squared error of an NMF model versus factorization rank for models of rank 1 to 100. An inflection point in the resulting curve was established around a rank of 35 (Figure 1a). A high-quality NMF model was fit within 1 minute with *k* = 35 using unnormalized counts and all genes. A k-Nearest Neighbor (k-NN) graph was constructed from sample loadings in this model and a UMAP embedding was learned to assess the sensitivity of the projection (Figure 1b). We observed generally distinct separation of cell types as annotated in the Human Embryonic Cell Atlas.

**Figure 1.**
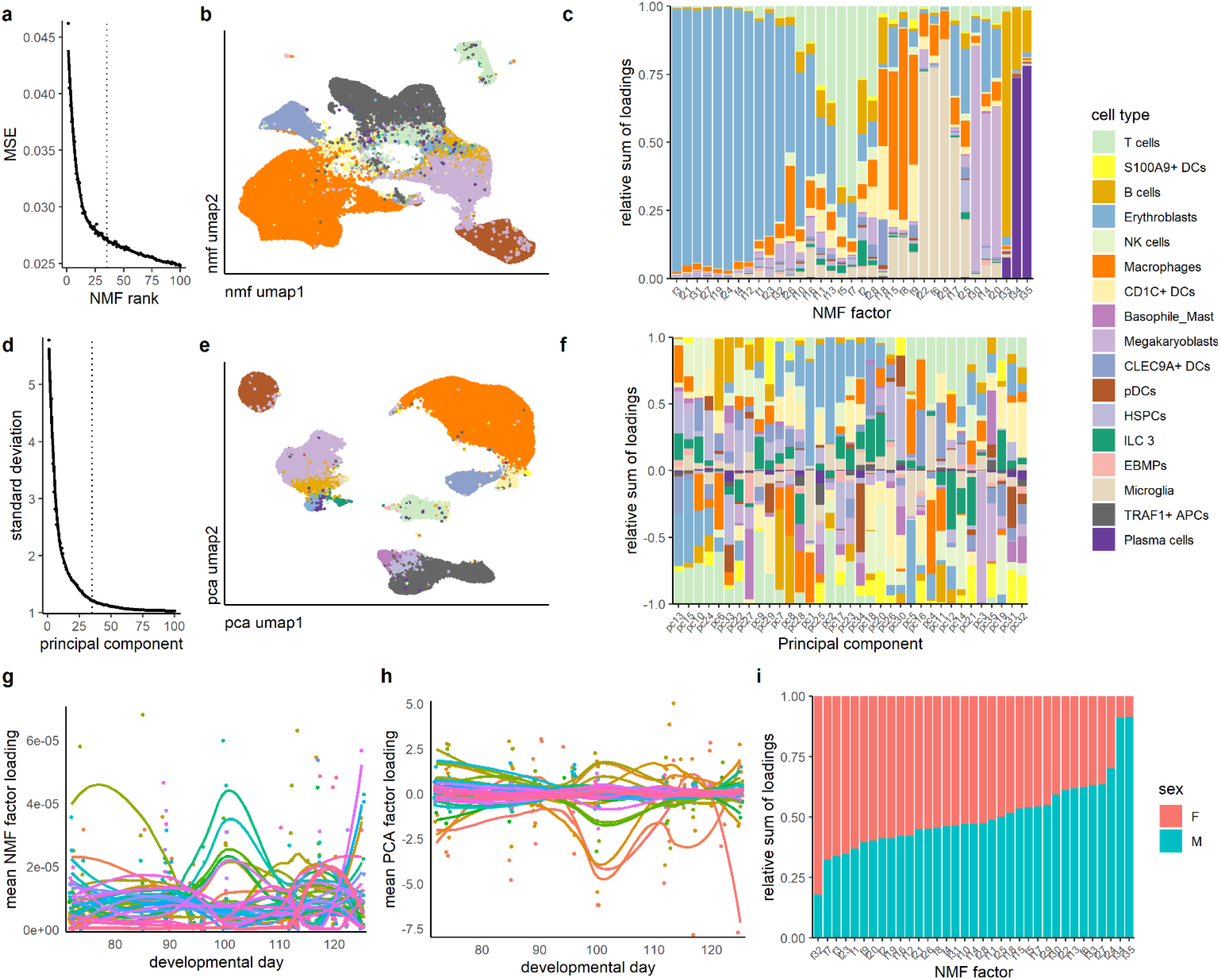
Dimensional reduction of 100,000 blood cells from the Human Embryonic Cell Atlas by NMF learns an interpretable and biologically relevant factor model, unlike PCA. a)Mean squared error (MSE) of NMF vs. factorization rank (*k* = 1 to 100). Model at *k* = 35 selected for analysis (dotted line). b)UMAP embedding from *k*-NN graph computed from sample loadings in NMF model, cells colored by cell type as annotated in the Human Embryonic Cell Atlas. c)Relative sum of loadings for all 35 NMF factors across all annotated cell types. d)Additional standard deviation explained by each principal component (1 to 100 PCs) from a PCA using the top 3000 highly variable genes. Model at *k* = 35 was selected for analysis (dotted line). e)As in (b) for PCA. f)As in (c) for PCA for all 35 principal components. g)Mean NMF factor loading in cells from embryos at various developmental days. h)As in (g) for PCA. i)Relative sum of factor loadings in all 35 NMF factors across cells from male or female embryos.

Principal component analysis (PCA) was run using the standard Seurat workflow. First, data was log-normalized (a step not needed in NMF), then scaled and centered (effectively converting a large sparse matrix to dense), then PCA was run for the top 3000 highly variable genes. The inflection point in the “elbow plot” of standard deviation vs. rank coincided with approximately *k* = 35 (Figure 1d). The resulting UMAP embedding clearly delineated atlas-annotated cell types (Figure 1e), perhaps better than NMF. The major caveat, however, is that cell types were originally classified with graph-based clustering on a PCA embedding by the atlas authors and thus are inherently biased towards PCA.

### NMF factors enrich for biological metadata relative to PCA

To examine whether the learned PCA embeddings encode biological information other than the rotation of coordinate axes to explain maximal variance, we examined whether factors in NMF or PCA were enriched for annotated cell types. We observed that some NMF factors were represented almost exclusively in certain cell types, others split between several cell types, and still others distributed across many cell types (Figure 1c). PCA factors, however, were randomly represented across all cell types (Figure 1f).

Samples from the Human Embryonic Cell Atlas span an 8-week window of development, enabling studies of differential gene expression over time. At least one NMF factor was specifically upregulated early around day 80, several factors were up around embryonic day 100, and still others were up late (Figure 1g). PCA factors manifested no such interpretable trends, with some factors down around developmental day 100, which is difficult to interpret (Figure 1h). This finding highlights the poor interpretability of signed decompositions of non-negative data.

NMF factors explain additional biological metadata as well. Some NMF factors were strongly enriched in cells from male embryos and at least one factor was strongly enriched in cells from female embryos (Figure 1i). There was also a strong association between some factors and cell source organ (Figure S2).

### NMF factors capture known biological processes more effectively than PCA

NMF factor models are additive decompositions of the input matrix. In the case of single-cell matrices of RNA transcript counts across cells, NMF factors should therefore represent biological processes (Clark et al., 2019; Stein-O’Brien et al., 2018). We examined GO term enrichment in NMF and SVD rank-35 models. SVD was used in place of PCA because PCA of 100,000 cells and 25,000 genes on a dense matrix takes too long, and no centering or scaling of a sparse matrix (conversion to dense) is necessary for SVD. Just over 3600 GO terms were significantly enriched in at least one of the SVD and NMF factors. The NMF factors manifested distinct patterns of GO term enrichment, indicating that factors describe largely unique biological processes (Figure 2a). However, SVD patterns manifested no such unique patterns, and factors that were enriched appeared to be non-specific (Figure 2b). Indeed, each NMF pattern had at least 100 enriched GO terms while most SVD patterns had none to very few (Figure 2c). These results show that SVD (and by extension, PCA) does not capture biological processes, but sequential normalizations or axes of associated variation, while NMF captures known genetic processes that collaborate to reconstruct transcriptional state.

**Figure 2.**
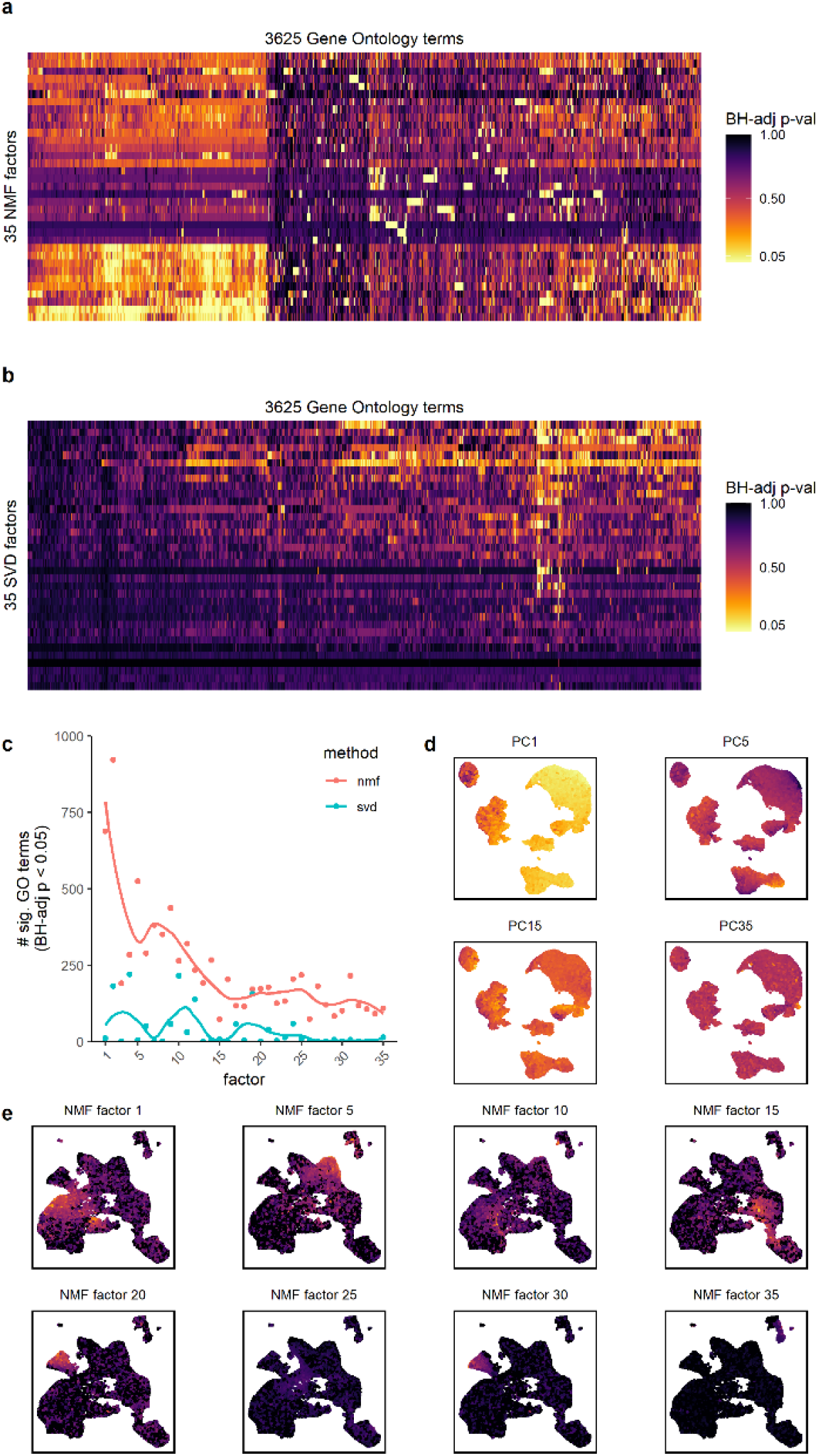
NMF factors describe biological processes that are differentially regulated across cell types, unlike SVD. a)Gene Ontology (msigdbr C5 human) term enrichment across all 35 NMF factors, BH-adjusted *p*-values are shown. All terms with at least one significant value in either the NMF or SVD gene set enrichment analysis are shown. Factors and terms are ordered by similarity. b)as in (a) but for the first 35 SVD factors. Factors and terms are ordered by similarity, not the same ordering as in (a). c)Number of significant GO terms (defined by BH-adjusted *p*-value < 0.05) in each NMF and SVD factor. Factors 1 to 35 are ranked by singular value (SVD) or proportion total signal explained (NMF). d)SVD sample loadings mapped to a UMAP embedding learned from the PCA (as in Figure 1E) for randomly selected components. e)NMF sample loadings mapped to a UMAP embedding learned from the NMF model (as in Figure 1b) for randomly selected factors. Color scale is a double square-root transformation for clarity.

We next examined how PCA embeddings and NMF loadings are dispersed throughout their respective UMAP embeddings. Principal component cell embeddings appeared as broad gradients passing through most cells, explaining a small fraction of the variation in that cell (Figure 2d). NMF loadings were highly specific, localizing to distinct populations of cells (Figure 2e). This again demonstrates that NMF factors describe basic biology in addition to serving as a sensitive dimensional reduction, thus doing what PCA is commonly used to do as well as providing additional information.

### RcppML NMF is as fast as state-of-the-art SVD

Widespread adoption of NMF as a method in single-cell analysis requires reasonable runtime. We benchmarked our implementation against NNLM NMF, the fastest implementation of simple NMF of which we are aware in the R community, and implicitly restarted Lanczos bidiagonalization SVD (irlba), a sparse matrix SVD known for its class-leading performance (Baglama & Reichel, 2005). For both 2,800 PBMCs (Figure 3a) and 40,000 bone marrow cells from the Human Cell Atlas (Figure 3b), RcppML NMF method performed nearly as well, or as well as irlba SVD across a range of ranks. Our method was more than an order of magnitude faster than NNLM NMF. RcppML NMF also performed impressively on benchmarking tests in random sparse matrices regardless of factorization rank, matrix sparsity, number of samples in the matrix, or rectangularity of the matrix (Figure S1).

**Figure 3.**
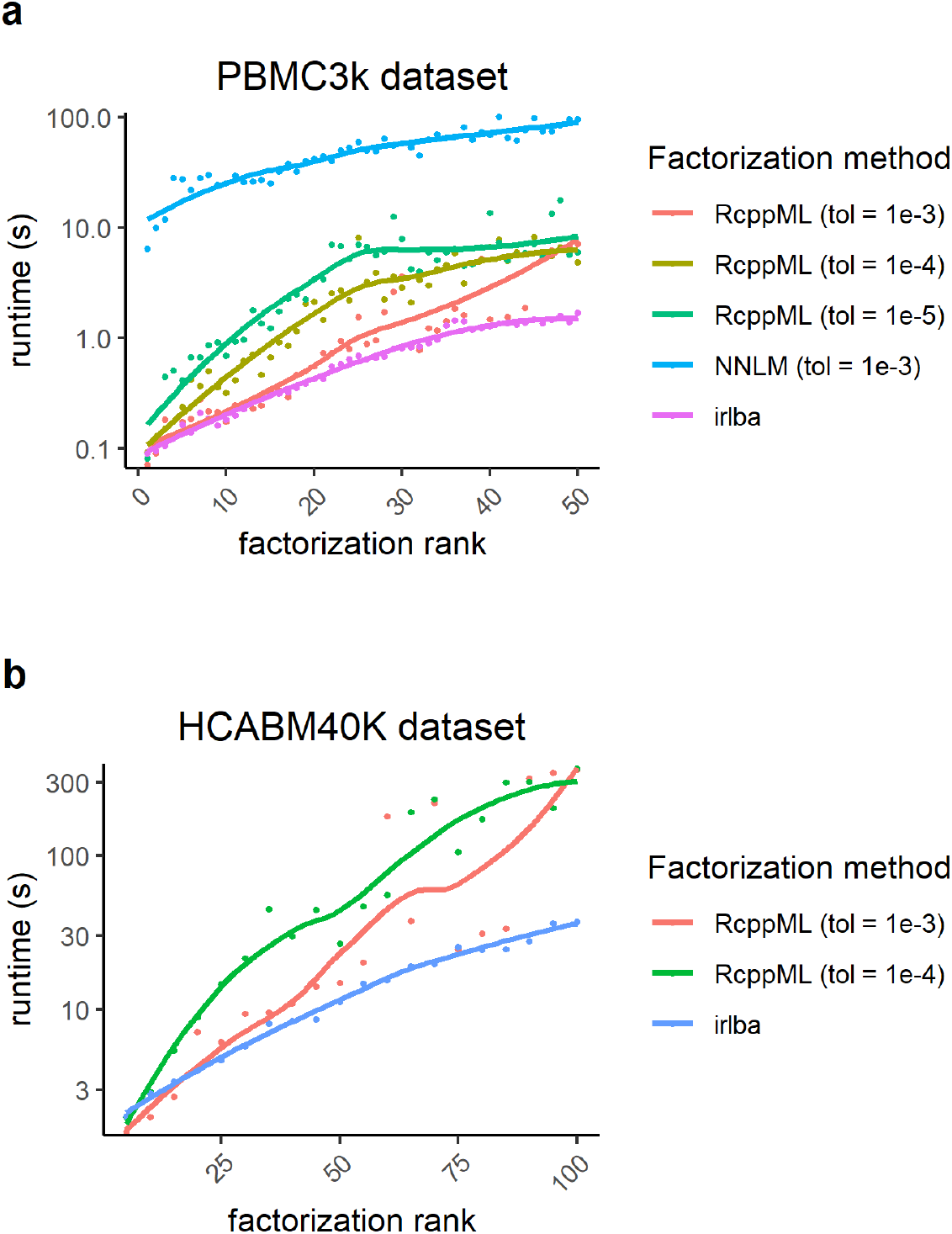
Runtime on an average desktop computer for RcppML NMF of single-cell datasets. a)Factorization of 2,800 cells from the PBMC3k dataset using irlba SVD, NNLM R package NMF, or RcppML NMF at two tolerances (1e-3 corresponds to cross-validation or rank-checking quality, 1e-4 corresponds to publication quality). Factorization ranks as indicated between 1 and 25. FAST-CD NNLS NMF without diagonalization was parallelized across 6 CPU threads. b)Factorization of 40,000 cells from the Human Cell Atlas Bone Marrow dataset (available through SeuratData). NMF settings as in (a).

### Efficient Non-Negative Least Squares solvers increase factorization speed

A major bottleneck in NMF is finding solutions to non-negative least squares equations. The NNLM R package has shown great promise for sequential coordinate descent least squares initialized with an approximate solution given by the model from the previous iteration (Lin & Boutros, 2020). We found that initializing coordinate descent non-negative least squares with a zero-filled vector created a strong gradient from which fast convergence could be achieved, faster than with warm-start initialization both with and without optimizers such as adam (see an example solution path in Figure 4a). We also considered initialization with an unconstrained least squares solution, or an approximate solution that we refer to as “Fast Active Set Tuning” (FAST). In FAST, an unconstrained least squares solution is computed, all negative values set to zero (an “active set”), and all other values are added to a “feasible set”. An unconstrained least squares solution is then solved for the “feasible set”, any negative values in the resulting solution are set to zero, and the process is repeated until the feasible set solution is strictly positive. The algorithm has a definite convergence guarantee because the feasible set will either converge or become smaller with each iteration. The result is generally exact or nearly exact for small well-conditioned systems (< 50 variables) within 2 iterations and thus quickly sets up coordinate descent very well (see example solution path in Figure 4b). The FAST method is similar to the first phase of the previously described so-called “TNT-NN” algorithm (Myre et al., 2017), but the latter half of that method relies heavily on heuristics to find the true active set, which we avoid by using coordinate descent instead.

**Figure 4.**
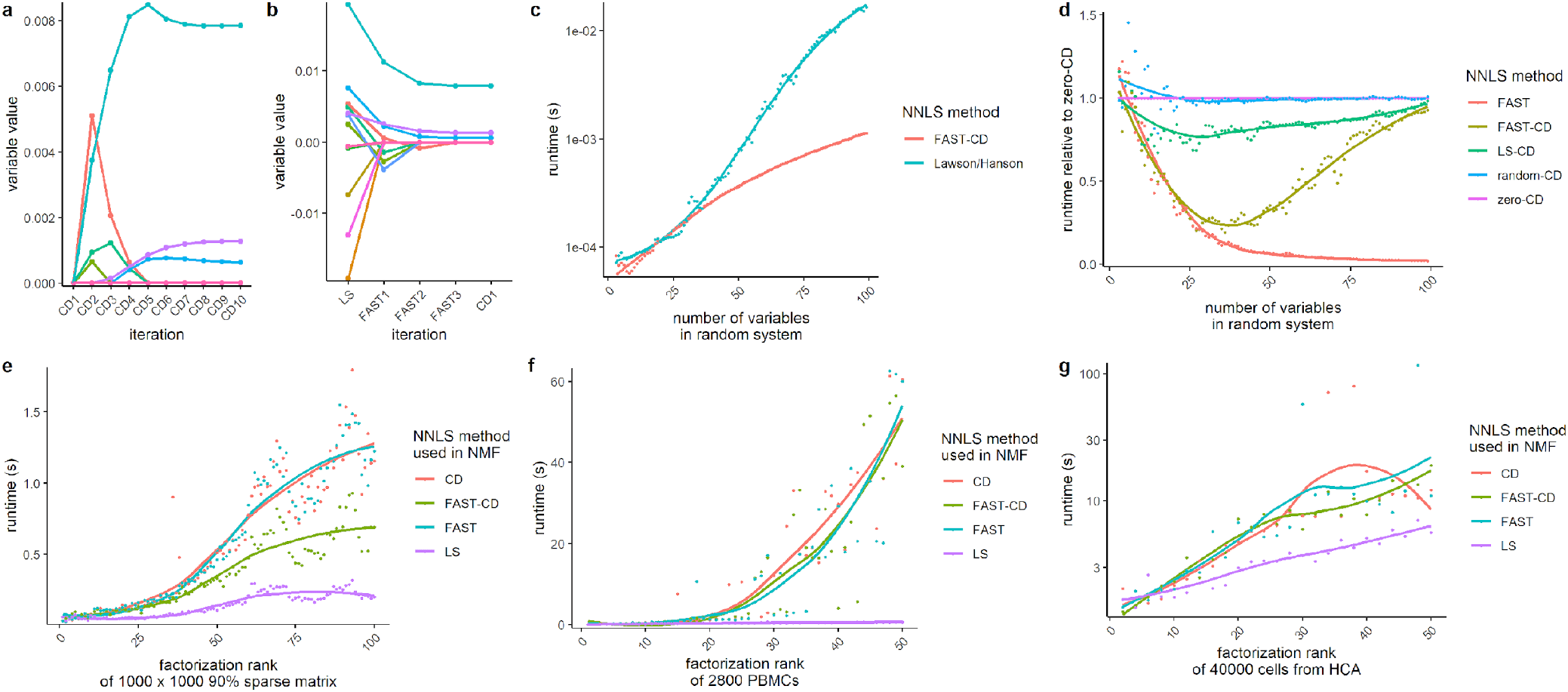
Novel non-negative least squares (NNLS) algorithms for fast NMF. a)Solution path for zero-initialized sequential coordinate descent least squares (CD) for a random ill-conditioned 12-variable system, where each variable is denoted by a color. 10 iterations were required for convergence to the actual solution. b)Solution path for Least Squares (LS), followed by Fast Active Set Tuning (FAST), followed by coordinate descent (CD) for the same ill-conditioned 12-variable system as in (A), where each variable is denoted by a color. 3 FAST iterations were needed to converge to the actual solution. A single CD iteration was needed to prove the actual solution was reached, and no refinement was necessary. c)Runtime of Fast Active Set Tuning (FAST) NNLS followed by refinement with coordinate descent (CD) compared to the gold-standard Lawson-Hanson fortran77 NNLS implementation as made available in the R function “nnls::nnls”. Benchmarks account for time required to compute cross-product *XtX* and *XtY*, as Lawson-Hanson computes *a = XtX* and *b = XtY* while RcppML::nnls simply solves *a* and *b*. d)Runtime of various least squares algorithms in solving ill-conditioned random systems between 2 and 100 variables in size: FAST (no refinement by coordinate descent, solutions may not be exact), FAST-CD (FAST-initialized coordinate descent), LS-CD (unconstrained least squares initializes coordinate descent), random-CD (uniform random distribution scaled to mean solution value used to initialize coordinate descent), zero-CD (zero-initialized coordinate descent). All runtimes are presented relative to zero-initialized coordinate descent. e)Runtime of RcppML NMF of a 1000 x 1000 90% sparse random matrix using least squares algorithms: CD (coordinate descent initialized with solution from previous iteration), FAST-CD (coordinate descent initialized with FAST approximation), FAST (FAST without coordinate descent), LS (unconstrained least squares, no non-negativity constraint applied). f)Runtime of RcppML NMF of 2800 cells from the PBMC3K dataset as in (e). g)Runtime of RcppML NMF of 40000 cells from the Human Cell Atlas Bone Marrow dataset (available through SeuratData) as in (e).

A fortran77 implementation of NNLS by Lawson-Hanson has long been the gold standard for NNLS, yet FAST-CD NNLS outperforms this standard for any random system greater than 30 variables in size (Figure 4c).

For solving random ill-conditioned systems, the FAST routine performs well indeed, but solutions deteriorate in quality as system size increases (Figure 4d). However, FAST-CD NNLS still outperforms coordinate descent NNLS initialized randomly, with zeros, or unconstrained least squares.

In the context of matrix factorization, zero-initialized coordinate descent performs better than FAST-CD on ill-conditioned random systems (Figure 4e) while factorization of actual single-cell datasets is approximately as fast, if not faster with FAST-CD compared to coordinate descent alone (Figure 4f-g). However, due to the better overall performance of zero-initialized coordinate descent across a large number of ranks for well-conditioned systems, and more recent optimizations to the logic within the coordinate descent NNLS solver, our current RcppML version uses only coordinate descent NNLS.

### Diagonalized NMF enables symmetric factorization

Random initializations for NMF can cause differences in the relative distributions of values in both sides of the model, *w* and *h*. These differences in turn cause factors to be scaled differently relative to one another than they would be if distribution of loadings in *w* and *h* were comparable. This can be demonstrated by factorization of a symmetric factorization (Figure 5a). However, when each factor in *w* and *h* is linearly scaled to sum to 1 against a diagonal, the distribution of *w* and *h* always exists on the same scale and convergence toward symmetry is guaranteed (Figure 5b). The scaling diagonal of NMF is different in nature to that of an SVD, as NMF factors are collectively and simultaneously optimized, while SVD factors (for example in rank-truncated SVD) are optimized sequentially and are dependent on preceding factors (Figure 5c).

**Figure 5.**
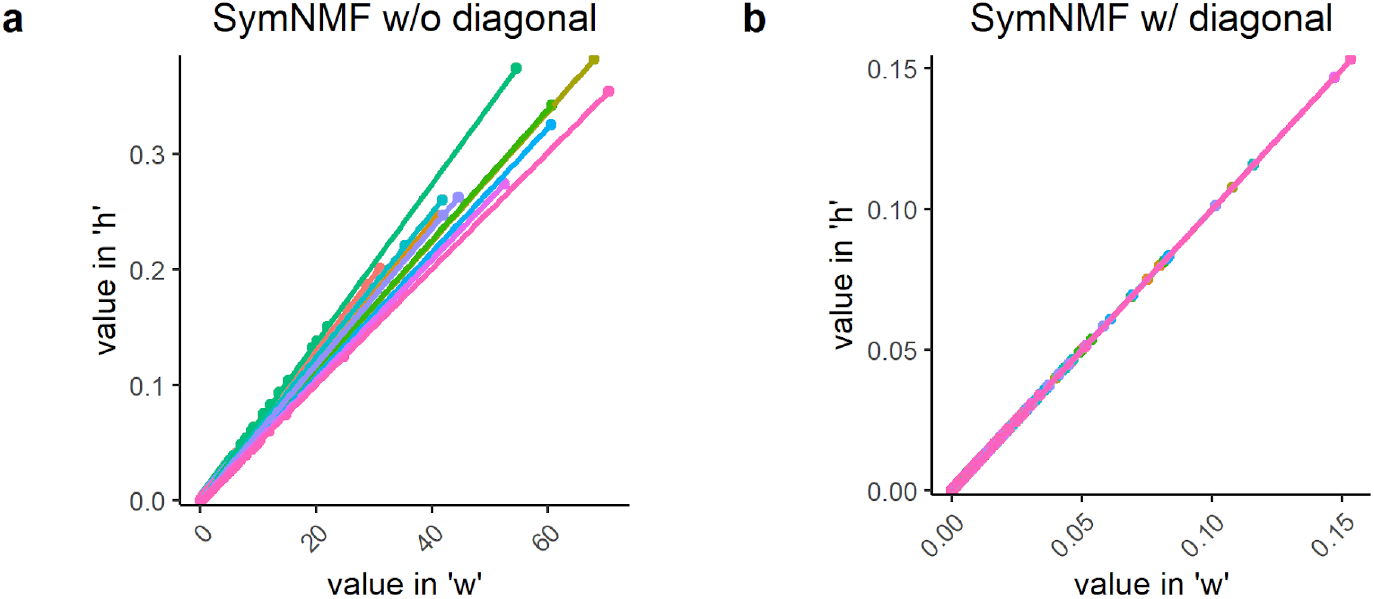
Diagonalization of NMF enables symmetric NMF and equal L1 regularization penalties across all factors. a)Rank-10 RcppML NMF of a symmetric 1000 x 1000 90% sparse matrix (SymNMF) of the form “*wh*”, thus without diagonal scaling. Loadings in “*w*” are plotted against corresponding loadings in “*h*”, colored by to which of the 10 factors they belong. b)Rank-10 RcppML NMF of the same matrix as in (a) of the form “*wdh*”, thus with diagonal scaling. Note that values in “*w*” and “*h*” are equivalent.

## Discussion

NMF is well-suited to dimensional reduction of single-cell data for visualization, use as an embedding for clustering, and learning patterns of coordinated gene activities. Our algorithm brings compute time down to that of the fastest SVD implementations and adds diagonalization to aid interpretability and convex regularization. Taken together, this work makes a case for adoption of NMF into mainstream single-cell analysis pipelines.

However, NMF shares some challenges with its signed dimensional reduction counterparts. For example, rank determination remains a subjective problem, although in Figure 1a we show that rank determination can be as “easy” as that for PCA using an inflection point in a curve of rank vs. model objective (loss for NMF and additional explained variance for PCA). Unfortunately, despite its intuitive appeal, this “elbow plot” method is highly subjective and often very unclear. Future work must explore more quantitative approaches such as cross-validation against robustness objectives or with missing value imputation (Lin & Boutros, 2020). Unfortunately, these approaches require the training of many models, and involve masking (which scales more poorly than the base algorithm), and themselves contain hyperparameters. However, rigorous rank-determination methods for PCA are at least equally consuming and ineffective as those for NMF, such as jackstraw (Chung and Storey 2015) or *k*-fold cross-validation. Rank determination for dimension reduction, at the end of the day, will remain a theoretical enigma and will always benefit from evaluation against domain knowledge and the art of expert inspection.

Reproducibility is yet another challenge for NMF. Alternating least squares requires an initialization, such as a random or non-negative double SVD model (Esposito, 2021). While NNDSVD is “robust”, it differs fundamentally in nature from NMF, and any non-random initialization can trap updates into a local minimum even if random noise is added to the model and zeros are filled. Different random initializations can lead to different solution paths toward local minima, as finding the global minima is NP-hard (Vavasis, 2009). Setting a random seed to guarantee reproducibility of a factorization model is far from ideal when the learned factors are intended to describe fundamental biology. However, for most applications and including scRNA-seq, replicate models give similar errors of reconstruction, many factors are robust, and those that are not as robust share the remaining information among themselves differently between replicates. Some heuristics have been proposed to find “consensus” NMF factors across multiple models (Kotliar et al., 2019; Stein-O’Brien et al., 2019), but these methods require significant compute time, no longer treat NMF as a dimensional reduction, and may be biased towards factors explaining dominant signals. While robustness may be seen as a weak link in NMF, it is important to realize that SVD is very sensitive to minor adversarial attacks on the input data. For example, different normalizations of the data or batch effects can lead to fundamentally different SVD results across most factors. On the other hand, because NMF factors are collectively updated, distinct technical issues are usually explained by a single factor while other factors are left unaffected. This same reasoning explains why no normalization of input data is required for NMF – the method naturally applies any necessary scalings or corrections.

NMF is an established method for extracting additive signals from non-negative data. For single-cell analysis, NMF describes coordinated gene activities and serves as a sensitive dimensional reduction for visualization and clustering. It seems more reasonable to cluster cells on factors describing biological processes than on principal components explaining variance. Further, it seems more reasonable to visualize cells based on underlying signals rather than embeddings learned from sequential normalizations of the data. Finally, while PCA and SVD are imputation-incompetent signed decompositions, NMF imputes missing values, denoises false positives, and respects non-negative inputs with non-negative outputs. Non-negative matrix factorization has the potential to systematically capture the complexities of genetic diversity, and it can do so simply, interpretably, and quickly.

## Materials and Methods

### NMF Problem Definition

Non-negative matrix factorization seeks to decompose a matrix A into orthogonal lower-rank non-negative matrices *w* and *h*:

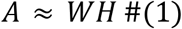

In the above equation and considering scRNA-seq datasets, *A* is a sparse matrix with unnormalized raw gene counts as rows and cells as columns, *w* is a dense matrix of “genes x factors”, and *h* is a dense matrix of “factors x cells”.

NMF algorithms generally require some initialization of *w* and/or *h* and use alternating updates of *w* and *h* to refine the model until convergence as determined by some stopping criteria. At convergence, *wh* will approximate the input *A*.

Alternatively, inspired by SVD or PCA, one might consider adding a scaling diagonal:

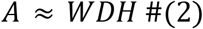

NMF may be diagonalized by scaling factors in *h* and *w* to sum to 1 after each alternating update. For example, after *h* is updated, *d* is set equal to the sum of factors in *h* and then each factor in *h* is scaled to sum to 1. Next, *w* is updated, after which *d* is set to the sum of factors in *w* and each factor in *w* is scaled to sum to 1.

### NMF Algorithm

RcppML NMF makes use of the traditional algorithm for NMF by alternating least squares (ALS) (Kim & Park, 2007). This algorithm is fast and flexible, especially compared to multiplicative updates (Lin & Boutros, 2020), even when supported by optimizers such as adam. In ALS-NMF, *w* and *h* are updated alternately with each iteration. For example, to update a column of *h* (*h*_*i*_) given a column of *A* (*A*_*i*_) and *w*, the Ordinary Least Squares (OLS) system is constructed as follows:

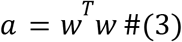

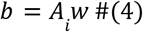

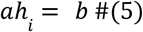

A similar approach may be used for updating rows of *w*, only *A* must first be transposed, and *h* used in place of *w* in the above equations. Equations #3-5 describe projection of a linear model which is a fundamental method in transfer learning. Projections of dense models onto sparse matrices may be performed using the R function “*RcppML::project*”.

### NMF Stopping Criteria

A well-established measure of convergence for matrix factorizations is given by the relative change in mean squared error (MSE) between consecutive iterations (Lin & Boutros, 2020). For a given iteration *i*, this measure is given by:

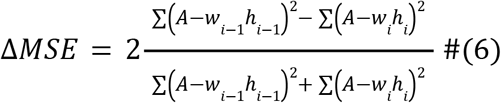

Calculating change in MSE requires computing the cross-product of *w* and *h* for each iteration to give a large dense approximation of the model in *wh*, then subtracting dense *wh* from sparse *A*. Such a dense-sparse operation, even with parallelization and proper use of sparse matrix iterators (as implemented in “*RcppML::mse*”), is extremely time-consuming, and in cases with >99% sparse matrices often takes longer than the factorization updates themselves.

Because MSE depends on *w* and *h*, the relative change in *w* and *h* across consecutive iterations may serve as a more efficient stopping criteria. Consider a stopping criterion using Pearson correlation (R^2^) between models in consecutive iterations. For a given iteration *i*:

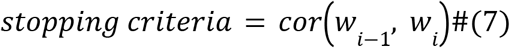

For a symmetric factorization, the stopping criteria could be correlation between *w* and *h*, since at convergence *w* ≈ *h*:

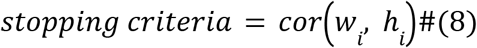

Indeed, for a symmetric factorization, the change in MSE between consecutive iterations, the correlation between w across consecutive iterations, and the correlation between w and h for a given iteration, follow extremely similar trends (Figure S3a). Since calculating correlation between models is at least an order of magnitude faster than calculating MSE (Figure S3b), RcppML NMF uses eqn. #7 as a stopping criterion in place of MSE.

### NMF Implementation

RcppML is a library of R functions that are lightweight wrappers around C++ functions coded with Rcpp and RcppEigen (Bates & Eddelbuettel, 2013; Eddelbuettel & François, 2011). For this scope of work, the Eigen C++ library is the fastest non-commercial BLAS and linear algebra library. We began by working in Armadillo, achieved comparable performance in base Rcpp, but found Eigen to be at least twice as fast for all use cases in addition to 3-5x faster at computing Cholesky decompositions. When plugging Intel MKL BLAS into Armadillo the gains for Eigen in our situation are still noticeable. All RcppML Eigen code is provided in a single C++ file so that functions can be readily incorporated into other C++ programs and software without using Rcpp wrappers, provided the Eigen header library is loaded.

An Rcpp sparse matrix class handles zero-copy pass-by-reference access to R objects in C++. This contrasts with Eigen and Armadillo sparse matrices, which form deep copies, and thus RcppML uses significantly less RAM. Constant sparse matrix forward column iterators are used for contiguous access to non-zero values. This provides performance gains for matrices >80% sparse, below which sparse iterators underperform dense operations. scRNA-seq data is typically 92-98% sparse and rarely <80%.

Block-pivoting in alternating updates of w and h require transposition of sparse matrix *A*. Transposition of very sparse column-major matrices is an inefficient operation. We profiled in-place updating of w and found that the computational cost of in-place updating surpassed the cost of up-front transposition after about 3 iterations. RcppML NMF therefore does not provide support for in-place updates of *w* and uses block-pivoting instead.

Parallelization with OpenMP is implemented across rows in *w* and columns in *h* for each alternating least squares update. By default, all available CPU threads are used. Calculation of the right-hand side of systems of equations is included in the parallelization loop alongside the least squares solver to minimize overhead and maximize the contiguity of memory access patterns within threads.

The FAST least squares algorithm relies heavily on the Eigen Cholesky module for extremely fast LLT decompositions and solutions by forward-backward substitution. For projecting a linear model, the LLT decomposition for the left-hand side of the system is computed only once (i.e. “preconditioned”). Sequential coordinate descent least squares is adapted from the NNLM RcppArmadillo package (Franc et al., 2005; Lin & Boutros, 2020), with significant optimizations to algorithm logic and vectorization.

Extensive code profiling and experimentation ensured we took full advantage of compile-time optimization and Eigen-facilitated vectorization. We were mindful of passing by reference, in-place updating, and contiguous memory access patterns for optimal use of the cache.

We have found that the base algorithm can be optimized for GPUs with CuSparse, but the gains were not significant enough to justify GPU over CPU utilization given the difference in cost. This is almost certainly due to the discontinuous cache access patterns in sparse-dense operations, which work well for CPU but not GPU operations. We have also explored multiplicative updates accelerated with the adam optimizer, which approached but did not surpass the runtime of our implementation. In the future we hope to add support for single precision, but beyond that we currently do not see further potential for speed gains. Our attention now turns to developing a neural network-like NMF implementation that can be extended much more flexibly to address problems beyond reconstruction such as integration, multimodality, generativity, and more.

### Benchmarks

All runtime benchmarks and nearly all figures were generated on an average desktop computer to show that any user can make good use of NMF for single-cell datasets. Unfortunately, for Figure 1 we had to use a high-performance computing cluster to run Seurat PCA and UMAP reductions due to RAM limitations. Benchmarks were run using a 64-bit OS running Windows 10 on an Intel Core i5-8400 CPU processor at 2.80GHz with 8GB of RAM. Runtimes were faster on a computing cluster, but relative performance between methods was similar. Random sparse matrices were generated using the “rsparsematrix” function from the R “Matrix” package. Runtimes for R function calls were measured using the “microbenchmark” R package.

### Dimension Reduction of Embryonic Blood Cells

NMF was run on standard log-transformed RNA transcript count data from 100,000 blood cells from the Human Embryonic Cell Atlas to 100 iterations at *k* = 35. PCA was run on log-normalized, centered, and scaled count data using Seurat default parameters. KNN graphs were constructed with *k* = 20 on dimensions 1 to 35, UMAP plots were constructed from KNN graphs using dimensions 1 to 35, 50 neighbors, and “Seurat::RunUMAP” defaults. Cellular metadata packaged with the Blood Cells dataset was used for enrichment analyses for cell type, organ type, embryo sex, and developmental day. GO enrichment analysis used msigdbr human C5 pathways, fast gene set enrichment analysis (fgsea), and BH-adjusted p-values. Many terms in the NMF factors were highly significant (*p*-adj < 1e-8), but the significance cutoff was defined at *p*-adj < 0.05 for convention.

## Data Availability

Figures 1 and 2 make use of the Human Embryonic Cell Atlas dataset, “Blood Cells” processed dataset available at https://descartes.brotmanbaty.org/bbi/human-gene-expression-during-development/, along with associated cell and gene metadata (Cao et al., 2020). Datasets used for benchmarking are readily available via SeuratData, consisting of the “pbmc3k” dataset of 2,800 PBMC cells supplied by 10x genomics, and the “hcabm40k” dataset, containing 40000 bone marrow cells from the Human Cell Atlas (Han et al., 2020).

## Code Availability

The latest stable release of the RcppML R package is publicly available on CRAN. The development version is available at github.com/zdebruine/RcppML. Issues on GitHub are monitored and we welcome questions and feature requests there.

